# Novel method for multiplexed full-length single-molecule sequencing of the human mitochondrial genome

**DOI:** 10.1101/2022.02.08.479581

**Authors:** Ieva Keraite, Philipp Becker, Davide Canevazzi, Maria C. Frias-López, Marc Dabad, Raúl Tonda-Hernandez, Ida Paramonov, Matthew John Ingham, Isabelle Brun-Heath, Jordi Leno, Anna Abulí, Elena Garcia-Arumí, Simon Heath, Marta Gut, Ivo Glynne Gut

## Abstract

Methods to reconstruct the mitochondrial DNA (mtDNA) sequence using short-read sequencing come with an inherent bias due to amplification and mapping. They can fail to determine the phase of variants, to capture multiple deletions and to cover the mitochondrial genome evenly. Long-read whole genome sequencing is prohibitively expensive for mtDNA heteroplasmy detection and often does not recapitulate the full mtDNA length.

Here we describe a method to target, multiplex and sequence full-length, native single-molecule the human mitochondrial genome utilizing the RNA-guided DNA endonuclease Cas9. Combining Cas9 induced breaks as barcodes with long-read sequencing, we implemented a protocol in an optimal setting for both high or low integrity genomic DNA to target the circular mitochondrial genome with extremely high coverage. Our analytical pipeline efficiently detects single nucleotide heteroplasmy, physically determines phase and can accurately disentangle complex deletion patterns. This workflow is a unique tool for studying mtDNA variation in health and disease, and will accelerate mitochondrial research.

## Introduction

Mitochondria are organelles found in most eukaryotic cells containing several copies of a circular genome encoding for vestigial functions of what once used to be a free-living organism. In human, in addition to the nuclear genome, cells have several hundreds to thousands of copies of the circular mitochondrial genome within mitochondria. With a reference sequence of 16,569 bases in size, the mtDNA encodes 13 proteins, 22 transfer RNAs, and two ribosomal RNAs^1–3^. Deep resequencing has uncovered a remarkable extent of mtDNA heteroplasmic variants in the human population^4^. The mtDNA with single nucleotide variants (SNVs) or with structural variants (SVs) can coexist with the wild-type mtDNA and can have substantial impact on human health depending on the variant position, frequency and other endogenous or exogenous factors. While low-level heteroplasmic mutations are likely functionally silent, higher mutation load may lead to severe clinical phenotypes^5,6^ and is associated with diseases affecting 1 in 4,300 individuals^4,7^. Hence, accurate detection and quantification is important for diagnosis, genetic counselling and treatment of mtDNA diseases.

Early techniques to detect heteroplasmy were based on mtDNA fragmentation with restriction enzymes^8–11^. Later more sensitive techniques were adapted that involved the PCR^11–16^. Although the detection limit of such techniques can reach ~0.01%, the targeted region in mtDNA is small and novel variations outside of it cannot be detected^14^. Massively parallel sequencing has revolutionized the analysis and the quantification of mtDNA variants. While the former gold standard, Sanger sequencing, has a detection limit of 10-15%, short-read sequencing methods perform with a precision >1% ^17,18^. These methods include pre-amplification by PCR, which itself adds polymerase errors and amplification related bias, and therefore they require careful interpretation^14,19–24^. Despite of its advantages, short-read sequencing is limited by the read length, which makes the determination of alternative variant phase impossible. Identifying multiple large-scale rearrangements is challenging but a requirement to achieve comprehensive mtDNA analysis^25^. In addition, mitochondrial DNA-like sequences, segments of mitochondrial DNA inserted into the nuclear genome throughout evolution and referred to as ‘nuclear mitochondrial sequences’ (NUMTs)^26,27^, can interfere in the bioinformatics analysis due to sequence similarity^28^. Therefore molecular and bioinformatics approaches are required to minimize the impact^29–31^.

Oxford Nanopore Technologies (ONT) and Pacific Biosciences (PacBio) are the only two companies marketing long-read sequencing capable of spanning most of the complex regions of the genome. One of the recent breakthrough enrichment techniques for long-reads is amplification-free targeted DNA sequencing using the Cas9 double-strand cutting enzyme. This method relies on preferential sequencing adaptor ligation to the cut sites of Cas9, but not the blocked ends of the uncut DNA molecules, therefore enriching for cleaved molecules. The method has been combined with the ONT^32^ and PacBio^33^ long-read sequencing platforms.

Here we present a method to target the mitochondrial genome selectively and sequence the entire native molecule in a single read starting from a Cas9 cut site, which serves, at the same time, as a multiplexing barcode to facilitate the workflow and decrease the cost per sample. The key step is opening the circular mtDNA molecules and making them accessible for library preparation. We have implemented and optimized the Cas9-mtDNA-enrichment protocol coupled with nanopore sequencing of multiple samples pooled on one ONT GridION flow cell. We also adapted our protocol to allow the analysis of low integrity genomic DNA (gDNA) samples. The combination of our method with the latest ONT Kit 12 Chemistry (Q20+ ligation sequencing kit) and the R10.4 flow cells, containing CsgG-family nanopores with an additional second constriction^34^, improves the overall read accuracy and the accuracy of homopolymer sequencing. Beyond the laboratory workflow, our developed informatics pipeline allows reliable detection of low-level heteroplasmies and disentangles complex deletion patterns. Translating this method into clinical research elevates our capacity to characterize the role of mtDNA in health and disease, opens a new avenue for population studies and can easily extend into other eukaryotes’ mitochondrial research.

## Results

### Preparing the Cas9-mtDNA-enrichment sequencing library

We adapted and further developed laboratory procedures, previously described for targeting nuclear DNA^32^, to highly selective single-molecule sequencing of mtDNA from a human gDNA sample. The method is amenable for multiplexing (Fig. 1) without the need for additional steps related to barcoding. To achieve efficient enrichment the method relies on the dephosphorylation of all available DNA 5′-ends in the gDNA sample, which is crucial to prevent ligation of nanopore sequencing adaptors to all non-targeted DNA fragments. After the nuclear DNA dephosphorylation step, a gDNA sample is split into aliquots and mtDNA in each aliquot is cleaved sequence-specifically by dual-RNA-guided Cas9 endonuclease (Fig. 1). This allows to selectively induce one double-strand break at a specific position in the mtDNA which is used as a barcode so that multiple sample aliquots, cleaved at different positions, can be joined for library preparation and sequenced in one sequencing flow cell without a need for additional indexes. The Cas9 cut-barcode also determines the full-length of the mtDNA as the molecule starts and ends with the same guide-directed Cas9 break. In our setting we used four guide RNA (gRNA) pairs, to multiplex four samples (Supplementary Table 1). This approach was modified for comprehensive characterisation of a clinical sample with multiple mtDNA deletions to a triplet of gRNAs. After Cas9 cleavage aliquots were treated with Proteinase K to remove the Cas9 protein bound to the mtDNA, pooled, and the ONT sequencing adaptors were ligated to the phosphorylated Cas9 cleavage sites. Method optimisation is provided in the Supplementary Material. For data analysis a dedicated informatic pipeline was developed.

**Fig. 1.**
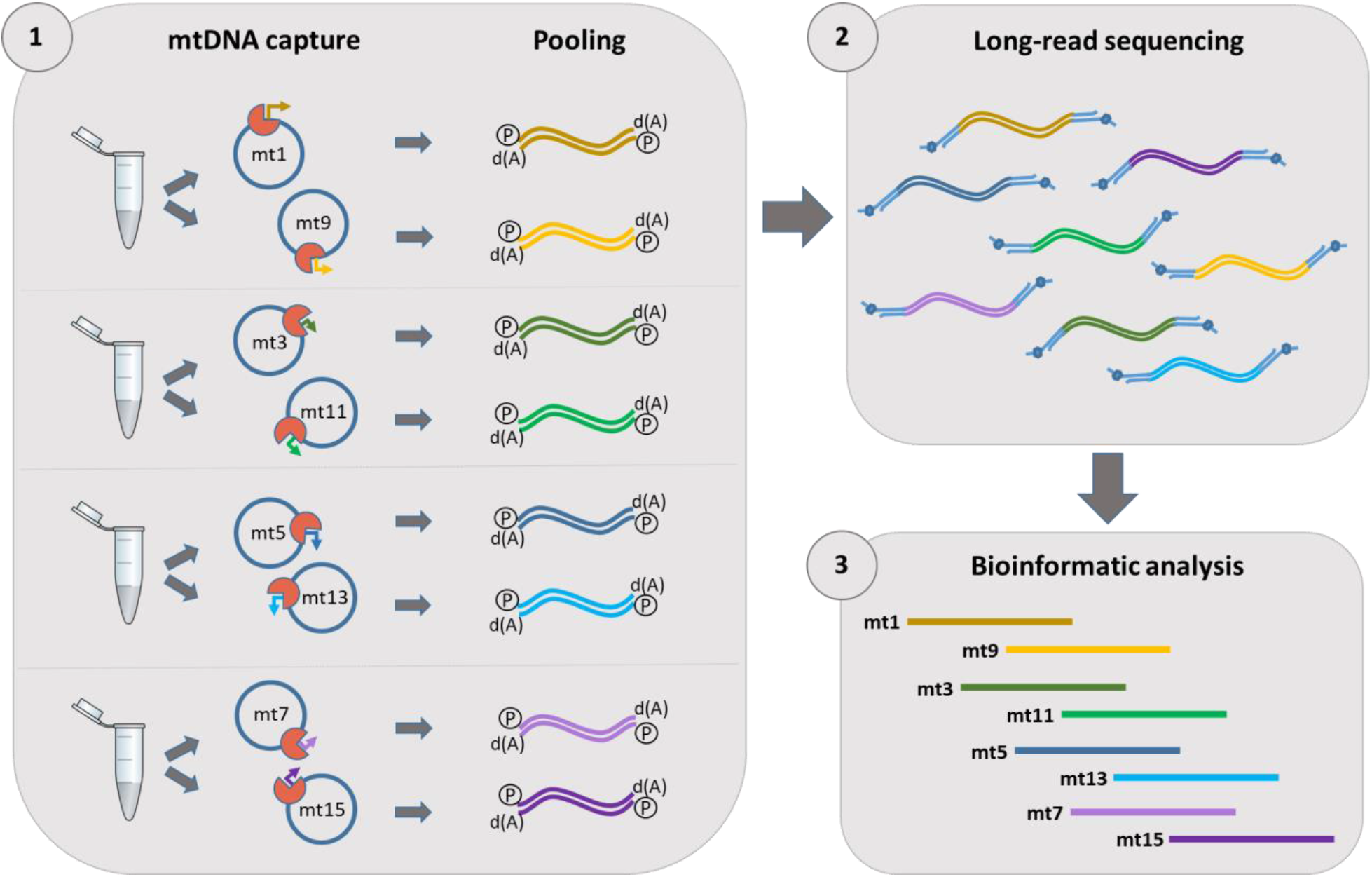
Cas9-mtDNA-enrichment, barcoding, pooling and demultiplexing approach for long-read sequencing. A schematic overview of the dual-guide targeting of full-length mtDNA with indicated gRNA-guided Cas9 cut sites. Samples can be targeted by several different gRNAs. Here we selected a pair of gRNAs on opposite sides of the circular mtDNA for each sample (1). After optional treatment of gDNA with Exonuclease V and dephosphorylation, samples are split into two aliquots for dual-guide targeted cleavage. Each cut site serves downstream as barcode in the analysis pipeline. The circular mtDNA molecules are opened, Cas9 is removed by Proteinase K digestion, which is followed by mtDNA dA-tailing and pooling of all of the aliquots. (2) ONT library preparation of the pooled samples and long-read sequencing on a nanopore flow cell. (3) The bioinformatics analysis encompasses basecalling, alignment, demultiplexing, variant calling and annotation.

### Demultiplexing

Demultiplexing of Cas9-mtDNA-enriched long-reads was performed by aligning the reads using minimap2 to the human whole genome reference GRCh38 to produce a PAF alignment file. Aligning mitochondrial reads can be problematic since alignment tools assume linear reference sequences, while the mitochondrial genome is circular. This can cause low levels of mapping of short sequence reads that span the transition between the end and start of the mitochondrial chromosome reference. With long-reads, however, this problem does not occur so a normal alignment procedure could be used for the demultiplexing step. However, a large fraction of the reads produced split or supplementary alignments where different segments of the read align to separate locations. This is expected for mitochondrial reads as any read that spans the end of the reference sequence will map as two segments, one ending at the end of the reference and the other starting at the beginning of the reference.

The PAF alignment records were used to select reads that uniquely aligned to the mitochondrial genome and, from these, to further select reads that started and/or ended near a cut site. The selected reads were also filtered to ensure that (a) all read segments aligned to the same strand of the mitochondrial genome; (b) the ordering of the segments within the read and along the chromosome were consistent (allowing for the circularity of the chromosome) and (c) almost all of the read could be successfully mapped to the mitochondrial reference sequence. Minimap2 can produce PAF (minimap2 specific) or a standard SAM output. PAF output was used for demultiplexing as it is a simpler format than SAM that made performing the previously described filtering easier.

Four different selection strategies for reads based on the mapping to cut sites were developed with different levels of stringency. The most stringent strategy, ‘**Both**’, requires that a read must both start and end near the same cut site. This has the effect of only selecting full-length reads (those that span the entire mitochondrial genome). The advantage of this strategy is that the risk of mis-assigning a read during demultiplexing is very low, the coverage across the mitochondrial reference is completely even in the absence of deletions, and the full-length reads do not jeopardize native read ratios due to length bias. If there are not enough full-length reads (for example, if the DNA sample is degraded) then less stringent strategies can be used. The ‘**Start**’ strategy only requires that a read starts near a cut site, while the ‘**Either**’ strategy will select a read if it starts or ends near a cut site. The less stringent strategies produce higher coverage but have the drawbacks of having an increased risk of mis-assignment of reads during the demultiplexing process, and also have uneven coverage across the mitochondrial genome. A fourth selection strategy was also tested purely for benchmarking and was designed to simulate the situation of dealing with highly degraded DNA when no full-length reads can be obtained. With this strategy, ‘**XOR**’, reads are selected either if the start or the end of the read matches a cut site, but not both. In this way, full-length reads are not selected with this strategy.

For all selection strategies, the matching of a read to a cut site was performed in the same way: the start (end) of a read matched a cut site if it lay not more than 100bp after (before) the cut site, taking the strand of the read into account when determining the direction. For example, a full-length read, selected with the ‘**Both**’ strategy, will either start at or just after a cut site and finish at or just before the same cut site if on the positive strand, or start at or just before a cut site and finish at or just after the same cut site if on the negative strand.

### Sequence variant calling, annotation and reporting

The results of the previous analysis stage were, for each selection strategy, a set of reads associated with each cut site. These reads then underwent a second alignment step, again using minimap2 but only to the mitochondrial genome reference and producing SAM output rather than PAF. The use of SAM output at this stage was to allow standard bioinformatics tools to be used for the downstream analyses. The alignment files belonging to each sample were merged using samtools merge before the subsequent steps.

SNV calls were made by producing a pileup of all reads assigned to a given sample. The consensus call and putative alternate allele were determined from the two most frequent bases seen at each site, while the alternate allele frequency (heteroplasmy) was estimated from the consensus and alternate base counts. Variant calls were filtered using a set of tests implemented in the program ont_align_view. This test set included: (a) a Fisher’s exact test of allelic strand bias; (b) a Fisher’s exact test of allelic bias in the number of bases filtered for low quality; (c) A t-test for a difference between average quality of the two alleles. Quality of heteroplasmic variants was estimated as the PHRED-scaled probability that the heteroplasmic frequency was 0. VCF files were produced with the called variants, and these were passed to the annotation pipeline. Variant annotations from the regularly updated resources dedicated to mtDNA were added to the VCF files: 1) disease-associated mutations and polymorphisms along with the GeneBank frequencies from MITOMAP^35^; 2) population frequencies for mtDNA variants from gnomAD^36^; 3) *in silico* pathogenicity prediction scores from MitoTIP^37^. In addition, mitochondrial haplogroup assignment was conducted for each sample with Haplocheck^38^. The complete variant annotation result output is summarized in the clinical samples variant annotation table (Supplementary Table 8).

SNV phasing was investigated from the pileup of all reads using the program ont_align_view. This program produces a text file with 16,569 columns, one for each position in the mitochondrial genome, and one row per read showing sequence bases, deletions and skips. For this file, the phasing of any set of positions (*e.g.*, a set of heteroplasmic positions) can be extracted and the frequencies obtained (Supplementary Fig. 2).

Deletion calls were made from examination of the sequence coverage. Regions of ≥ 5bp showing a significant reduction in coverage were identified. Each putative deletion was checked to verify that (a) it could be seen on both strands and (b) it was not in the region of a cut site.

### Cas9-mtDNA-enrichment to identify heteroplasmy in pathogenic SNVs

We applied our enrichment method, without the Exonuclease V treatment, on a set of 15 clinical samples, with a confirmed heteroplasmy of pathogenic SNVs established by a diagnostic laboratory (Table 1). The gDNA samples originated from different clinical specimens (Supplementary Table 2) and consequently the extracted DNA had variable quantity and integrity. The mtDNA genome coverage was between 33× and 2,335×, where the lowest coverage was in samples from oral mucosa and urine, consistent with the DNA degradation level determined from the DNA quality control (data not shown). We ran each low integrity sample in a separate sequencing flow cell, and in the analysis we included the full reads (**‘Both’** in Table 1) and the reads that started at the specific barcode (Cas9 introduced cut site) but were not exclusively full-length (**‘Start’** in Table 1). This strategy helped to increase the coverage 48–78-fold for these degraded samples and increased confidence of variant calling. The blood extracted DNA samples were less degraded, thus this strategy provided a less significant boost, mostly 2–3-fold, also these samples could be confidently multiplexed (demultiplexing data is provided in Supplementary Table 3). The identified SNVs with variant calling threshold cut-off ≥1% were concordant with the diagnostic laboratory results for all samples. One sample – AW6500 – that carried an extremely low heteroplasmic variant (<1%) had to be included in the annotation manually. We could confirm the heteroplasmy at the m.3243A>G position, which correlated with the report from the diagnostic laboratory (Table 1). No deletions were observed by our Cas9-mtDNA-enrichment or the lrPCR quality control (Supplementary Fig. 3 – Samples (2)). In addition, non-pathogenic polymorphic variants were identified in all samples mostly in homoplasmy or high-level heteroplasmy (>90%) along with variants of unknown significance which were observed mostly in low heteroplasmic frequency (<10%) (Supplementary Table 8). Assessing variant frequency within the general population and specific haplogroups is helpful when evaluating the pathogenicity of such variants^39^.

**Table 1.**
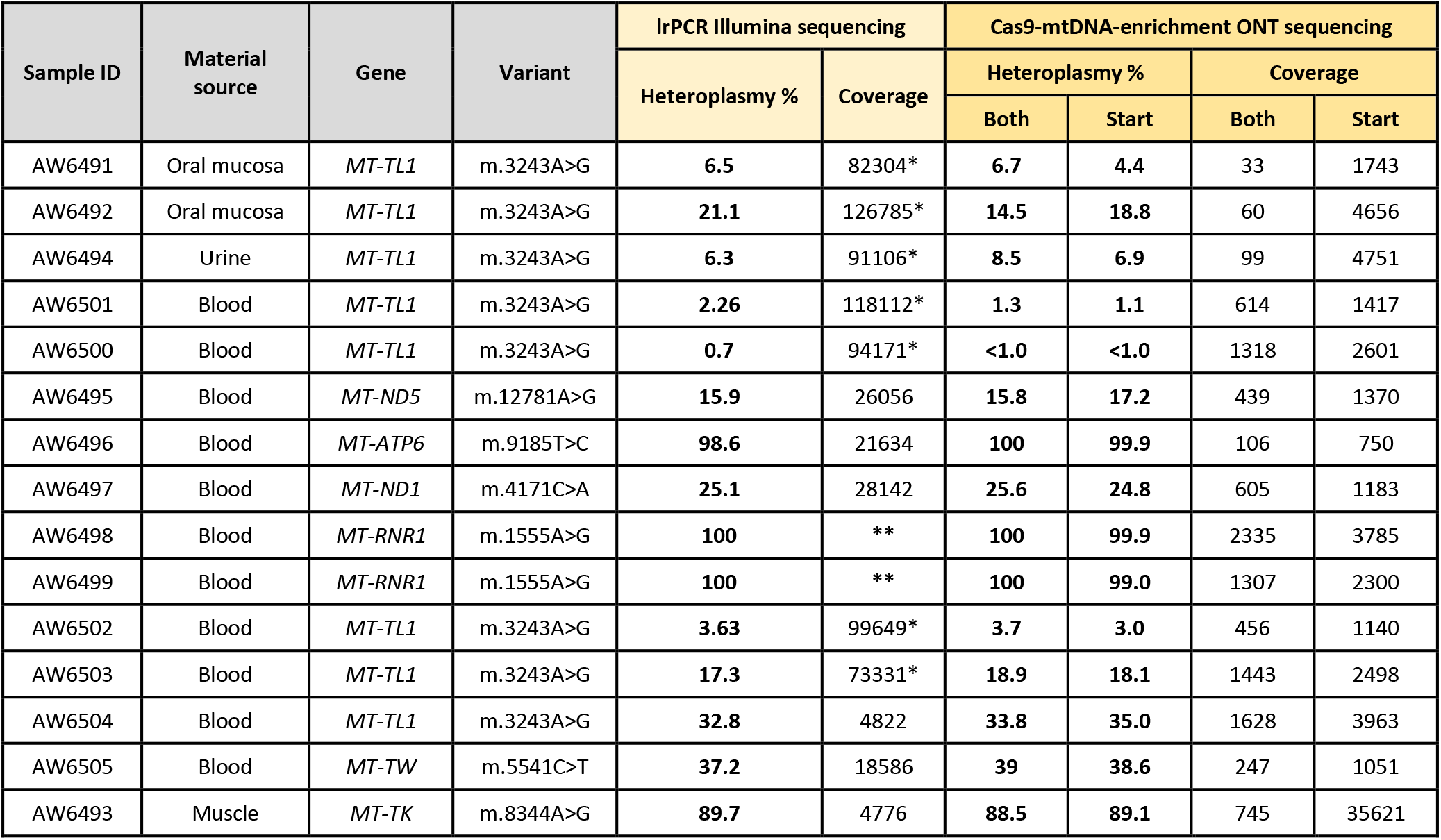
Pathogenic SNV identification and heteroplasmy determination in the confirmed clinical samples with mtDNA alterations.

The pathogenic variants identified by the Cas9-mtDNA-enrichment nanopore sequencing were compared with the clinical laboratory results. The low integrity gDNA samples display low coverage in the “**Both**” bioinformatics analysis, while the “**Start**” strategy allows boosting coverage. The determined heteroplasmies in all clinical samples are concordant with the diagnostic laboratory results. * Full-length mtDNA was not studied, only m.3243 region was amplified; ** Sanger sequencing.

### Application of the Cas9-mtDNA-enrichment to identify multiple deletions

To scrutinize our method to detect multiple mtDNA deletions, we used a gDNA (AW6506) sample extracted from a muscle biopsy of an entangled clinical case known to have complex mtDNA SVs. We analysed the gDNA sample without prior information about the positions of potential deletions. In this case, the diagnostic laboratory had been able to identify the exact position only for a smaller deletion, and an approximate position of a larger deletion, using lrPCR and Illumina sequencing. The copy number of these structural variants was not quantified precisely due to too high PCR biases.

In our quality control by lrPCR, we observed two bands representing two mtDNA populations of ~13.5 kb and ~3.5 kb in this sample. Wild-type 16.5 kb mtDNA was not visible on the gel, due to the amplification bias against larger fragments (Supplementary Fig. 3 – Sample (3)). The presence of the wild-type mtDNA molecule and both deletions was confirmed by ONT and Illumina sequencing of the lrPCR product, however, the sequencing results were heavily disproportioned by the amplification bias (Fig. 2a, b). Considering the limited gDNA sample availability and the complexity of the analysis, we used three gRNAs to identify the different SVs and ran this sample on a GridIon sequencing flow cell on its own. We selected one gRNA mt3 targeting at position m.3127 near the lrPCR primers annealing site, and another two gRNAs, mt5 and mt11 at mtDNA positions m.5142 and m.11239, respectively, to span the mitochondrial genome (Fig. 2c–e). We successfully detected two deletions present in the sample. A large deletion m.3257_16071del, reported previously by others^40^, was detected at a low-level heteroplasmy (8.1%) covering a segment encoding 13 polypeptides crucial for the oxidative phosphorylation. A small deletion m.10751_14129del containing 3 essential polypeptides^41^ was detected in 84.4% of reads. The wild-type mtDNA was present in 7.4% of reads. By comparing the nanopore sequencing results from the lrPCR amplicons and Cas9-mtDNA-enrichment, we exemplify the benefit of the native mtDNA sequencing. We observed the significant bias of preferential short molecule build-up resulting in inaccurate estimates of the three mtDNA populations in the lrPCR nanopore sequencing (Fig. 2f).

**Fig. 2.**
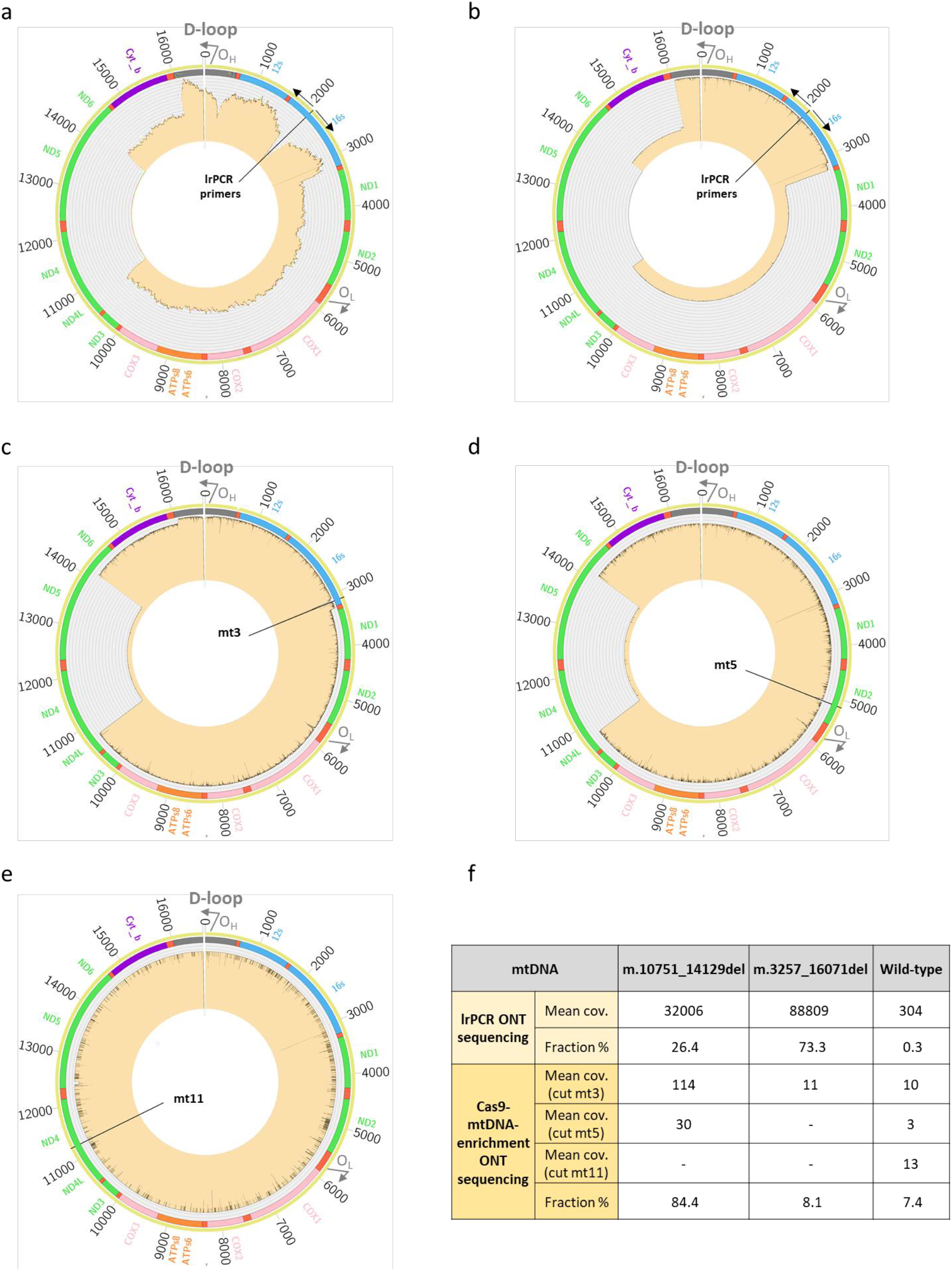
Multiple mtDNA deletions in a clinical sample. **a**, **b**, Circos plot of the lrPCR products of sample AW6506 showing three full-length lrPCR amplicons – two deletions and wild-type, sequenced by Illumina short-read (**a**) and ONT long-read (**b**) instruments. Black arrows at positions m.2120 and m.2119 represent forward and reverse primers, respectively. **c**, **d**, **e**, Circos plots of sample AW6506 targeted with the Cas9-mtDNA-enrichment sequenced on a GridIon flow cell showing three populations of mtDNA. **c**, Targeted with the mt3 gRNA at m.3127, all three populations can be observed – two deletions and the wild-type; **d**, mt5 gRNA at m.5142 results in two populations – the small deletion and the wild-type; **e**, mt11 gRNA at m.11239 results in capturing the wild-type population only. In the circos plots, the reference circle colors denote genes encoding protein subunits of complex I (green), III (purple), IV (pink), V (orange), ribosomal RNAs (blue), transfer RNAs (red), and non-coding region D-loop (grey). **f**, Table summarizing the comparison of mtDNA population fractions in lrPCR amplicons and Cas9-mtDNA-enriched native molecules, both sequenced on GridIon. Mean coverage of selected full-length reads and SV proportion are provided.

## Discussion

Here we present for the first time an efficient and selective targeting method for the mitochondrial genome delineating its native full-length sequence status using long-read sequencing. Due to lack of unbiased genomic tools to study the mtDNA we repurposed nanopore Cas9-targeted sequencing (nCATS)^32^ and with an effective optimization used Cas9 RNA-guided nuclease to target the circular extranuclear human DNA located in mitochondria. We achieved high enrichment and even coverage of the mitochondrial genome using full-length native reads, identified by the RNA-guided Cas9 cut site. Different cut sites were used to multiplex and pool different samples. The method was scrutinized for the effectiveness of multiplexing and precise targeting of full-length mtDNA molecules followed by sensitive detection of SNVs or disentangling complex multiple deletions in 16 clinical samples. Our study revealed undeniable benefits of applying the Cas9-mtDNA-enrichment protocol and nanopore native sequencing, exemplified by the distorted lrPCR amplicon representation. Although the basic strategy for variant calling of the mitochondrial genome is the same as for any genomic sequencing study, the challenges of handling the circular genome and the fact that the heteroplasmic variants can have any frequency requires a custom analysis. While variant calling pipelines for mtDNA exist (for example, GATK mitochondrial calling pipeline), the strategies used to handle the circular genome are not applicable to long sequence reads that can span the full length of the mitochondrial reference genome. It was therefore necessary to develop a custom bioinformatics analytical pipeline that can take advantage of the unorthodox biological characteristics of the mitochondrial genome and the particularities of long-reads from the nanopore sequencing platform to produce an efficient automated pipeline for the demultiplexing, variant calling, filtering and annotation. The pipeline attains a sensitivity of 98% using the R10.4 flow cells with Kit 12 Chemistry and a specificity >99.9%, and also allows accurate estimation of SNVs phasing frequencies. In addition, our system allows for complex rearrangements to be detected and characterized with accurate estimation of breakpoints and frequencies.

Currently, the detection of the mitochondrial sequence variants is a multistep process involving several molecular methods^25^. Designed to enrich the full-length mitochondrial genome in a single test our method offers a cost-effective native mtDNA sequence analysis with high coverage for accurate detection of SNVs, to define accurately single or multiple deletions, and to reconstruct haplotype lineages. Together with its analytical pipeline it is a promising tool for translational research to fill a void in current approaches to molecular diagnosis of mtDNA-related disorders and provides unique leverage to mtDNA population studies, including the confounding biases related to NUMTs. Cas9-mtDNA-enrichment could potentially be used for the investigation of somatic mutation events in the mitochondrial genome. Directly sequencing the native DNA that is captured by the Cas9-mtDNA-enrichment also opens the door to investigate any DNA modifications the mtDNA might harbour. The protocol will likely enable extending the subject beyond human mtDNA to other eukaryotes where mitochondrial genome size can be an impediment for lrPCR full-length amplification.

## Materials and methods

### Human cell lines

Cell lines HEK293, A549, CAPAN-2, SH-SY5Y were originally obtained from ATCC. Cells were cultured according to recommended protocols and were maintained at 37°C in 5% CO_2_. DNA was extracted from pellets containing 2-3 million cells using the Nanobind CBB Big DNA kit (Circulomics) and stored at 4°C until use.

### Clinical samples

A set of 16 clinical samples was previously collected. The DNA extraction was performed at Hospital Universitari Vall d’Hebron (Supplementary Table 2) from blood samples using chemagic DNA Blood400 Kit LH (Perkin Elmer). The DNA from oral mucosa swabs, urine and tissue samples was extracted using Gentra Puregene Blood Kit (Qiagen). All subjects gave informed consent approved by the local bioethics committee and in accordance with the Declaration of Helsinki. For the purpose of clinical diagnosis the whole mitochondrial genome was amplified in a single amplicon by lrPCR using the Takara LA PCR kit as previously described^42^ using 10 ng of DNA per reaction. Qubit 2.0 Fluorometer (Thermo Fisher Scientific) was used to quantify the amplicons and each sample normalized to 0.2 ng/μl. Amplicons were fragmented using NEBNext dsDNA Fragmentase (NEB), and a library was prepared using the NEBNext Ultra II DNA library prep kit for Illumina (NEB) following the manufacturer’s instructions. The pooled, indexed libraries were loaded into the MiSeq Reagent kit V2, 300 cycles (Illumina) and sequenced on the MiSeq platform (Illumina) in paired-end mode with a read length 2×151bp. In general, 40 libraries are multiplexed to obtain 5,000× mean coverage. In the mtDNA pipeline, quality filtering and trimming of overrepresented sequences was done using Trimmomatic^43^ to eliminate the sequences corresponding to the cut adapters used. After the trimming, the reads were mapped against the GRCh38 reference genome using BWA-MEM^44^. For variant calling and annotation we used Pisces^45^ with the following parameters: Depth ≥ 10, variant q-score between 20 and 100, variant frequency ≥ 0.01 and base quality ≥ 20.

### DNA quality control

DNA was quantified using a Qubit fluorometer with the dsDNA Broad Range Assay kit (Thermo Fisher Scientific) following the manufacturer’s instructions. DNA purity was evaluated using Nanodrop 2000 (Thermo Fisher Scientific) UV/Vis measurements. To determine the cell line gDNA integrity pulse-field gel electrophoresis, using the Pippin Pulse (Sage Science) was performed. For this analysis a SeaKem® GOLD Agarose 1% (Lonza) gel was prepared in 0.5× TBE buffer (Thermo Fisher Scientific). Approximately 150 ng of DNA sample was loaded together with CHEF DNA Size Standard Ladder (BIO-RAD) and Quick-Load 1 kb Extend DNA Ladder (NEB) to aid size determination. Fragments were separated using a pre-set 5-80 kb protocol. After the run, the gel was visualized using a NuGenius imaging system (Syngene). DNA integrity of the clinical samples was assessed with the Femto Pulse System using the Genomic DNA 165 kb Kit (Agilent Technologies) following the manufacturer’s protocol. The gDNA samples were stored at 4°C.

### Long-range PCR (lrPCR)

Full-length mtDNA amplification was carried out for short-read and long-read sequencing, and as a part of quality control using Platinum SuperFi II DNA Polymerase (Invitrogen). A forward primer Mt2120F (5’-GGA CAC TAG GAA AAA ACC TTG TAG AGA GAG −3’) and a reverse primer Mt2119R (5’-AAA GAG CTG TTC CTC TTT GGA CTA ACA −3’) were used. All lrPCRs (50 μl) were prepared as follows: 1× SuperFi II buffer, 0.4 mM each of dNTPs, 0.2 μM of each forward and reverse primers, 1× of SuperFi II DNA Polymerase, 0.5 mM MgCl_2_, and 10–100 ng of individual DNA sample. Cycling conditions were: 94°C for 1 min; 30 cycles of 98°C for 10 s, 68°C for 16 min; 72°C for 10 min; hold at 4°C. To visualize amplicons, Ultra Pure Agarose 1% (Invitrogen) gel was prepared in 1× TAE buffer (BIO-RAD). DNA samples were loaded together with Quick-Load 1 kb Extend DNA Ladder (NEB) to aid size determination. Fragments were separated running a gel at 60 V for 2.5-3 h. Amplicons were purified with AMPure XP beads (Agencourt, Beckman Coulter) and eluted in 30 μl nuclease-free water. Concentration was measured using a Qubit fluorometer with the dsDNA Broad Range Assay kit (Thermo Fisher Scientific) according to the manufacturer’s instructions.

### Preparation of Cas9 ribonucleoprotein complexes (RNPs)

To perform a Cas9-mtDNA-enrichment of full-length mtDNA sequences on single or multiplexed up to 4 samples per flow cell, a modified Cas9 protocol was established. A respective number of crRNA:tracrRNA duplex pairs were prepared. Here, 10 μM of every crRNA in TE pH 7.5, (IDT) was annealed with 10 μM of tracrRNA (IDT) separately in Nuclease-free Duplex Buffer (IDT) by denaturation at 95°C for 5 min and cooling down to RT for 10 min. Then, 10 μM of each crRNA:tracrRNA duplex was assembled together with 0.5 μM of Alt-R® S.p. HiFi Cas9 Nuclease V3 (IDT), 1× CutSmart Buffer (NEB) in nuclease-free water to form RNPs. This mix was incubated at RT for 30 min. After incubation, formed RNPs were held at 4°C until further use.

### Circular mtDNA pre-enrichment

To digest linear DNA 1 μg of DNA sample was incubated in 50 μl with 1× NEBuffer 4 (NEB), 1mM ATP (NEB) and 10 units of Exonuclease V (NEB) in nuclease-free water. The reaction was incubated at 37°C for 30 min and heat inactivated at 70°C for 30 min. Each treated DNA sample was cleaned-up with AMPure XP beads (Agencourt, Beckman Coulter) and eluted in 24 μl nuclease-free water.

### gDNA dephosphorylation

The gDNA dephosphorylation was carried out in a total volume of 30 μl containing 1 μg DNA, 1× CutSmart Buffer (NEB) and 15 units of Quick CIP (NEB) by incubating at 37°C for 20 min followed by Quick CIP inactivation at 80°C for 3 min. For the following Cas9-mtDNA-enrichment procedure each DNA sample was split into aliquots.

### Cas9-mtDNA-enrichment, library preparation, multiplexing and long-read sequencing

Each dephosphorylated DNA sample aliquot (15 μl) was mixed with 5 μl of appropriate RNP complex, and incubated for 20 min at 37°C. The mtDNA digestion was followed by incubation with 0.12 units of Thermolabile Proteinase K treatment (NEB) for 15 min at 37°C and 10 min at 55°C for inactivation. The dA-tailing was done by adding 0.25 mM dATP (NEB), and 0.5 μl Taq polymerase (NEB) for 10 min incubation at 72°C. Following dA tailing, all processed DNA aliquots were pooled together. The DNA sample pool was purified with AMPure XP beads (Agencourt, Beckman Coulter) and eluted in 42 μl nuclease-free water and processed with ligation sequencing Kit 12 Chemistry, SQK-LSK112 (ONT). The ligation mix was prepared in 38 μl containing 20 μl of Ligation Buffer (ONT), 5 μl of Adapter Mix H (ONT), 10 μl of T4 DNA ligase (NEB), and 3 μl of nuclease-free water. The ligation mix with the library was incubated for 1 h at RT. Ligation was terminated with TE buffer, pH 8, sample incubated with AMPure XP beads (Agencourt, Beckman Coulter) and washed twice with short fragment buffer (ONT). The sequencing flow cell was prepared according to ONT recommendations and Loading Beads II (ONT) were added to the library immediately prior to loading of the sample.

The library was sequenced on GridION Mk1 using the R9.4, R10.3 or R10.4 flow cells (ONT). Prior to every experiment, quality control was performed for each GridION flow cell using the MinKNOW software (v.21.05.20-21.10.8). Sequencing experiments were running for 72 h with off-line basecalling.

### lrPCR amplicons long-read nanopore sequencing

Library preparation was started with DNA repair and end-prep of 50 fmols of lrPCR mtDNA amplification product using NEBNext FFPE DNA Repair Buffer (NEB), NEBNext FFPE DNA Repair Mix (NEB), Ultra II End-prep reaction buffer (NEB) and Ultra II End-prep enzyme mix (NEB) incubated at 20°C for 30 min and at 65°C for 30 min according to ONT protocol. Each sample was purified with AMPure XP beads (Agencourt, Beckman Coulter) and eluted in 60 μl nuclease-free water. For adapter ligation, elution was mixed with 25 μl Ligation Buffer (ONT), 10 μl NEBNext Quick T4 DNA Ligase (NEB) and 5 μl Adapter Mix H (ONT). The reaction was incubated for 1 h at RT. Ligation was terminated with a clean-up using 0.4× volume of AMPure XP beads (Agencourt, Beckman Coulter). The library was quantified using a Qubit fluorometer with the dsDNA Broad Range Assay kit (Thermo Fisher Scientific). Approximately 10 fmol of library was loaded on the R10.4 flow cell and sequenced on GridION Mk1 following manufacturer’s recommendations. Sequencing experiment was set to run for 100 h with off-line basecalling.

### lrPCR amplicons short-read sequencing

Short-insert paired-end libraries of the lrPCR mtDNA product of the cell lines were prepared with KAPA HyperPrep kit (Roche) with some modifications. In short, 1 μg of lrPCR DNA was sheared on a Covaris™ LE220-Plus (Covaris). The fragmented DNA was end-repaired, adenylated and Illumina platform compatible adaptors with unique dual indexes and unique molecular identifiers (IDT) were ligated. The libraries were quality controlled on an Agilent 2100 Bioanalyzer with the DNA 7500 assay for size and the concentration was estimated using quantitative PCR with the KAPA Library Quantification Kit Illumina Platforms (Roche).

The library preparation from the lrPCR product originating from the clinical samples was carried out with Illumina DNA PCR-Free Library Prep, Tagmentation Kit (Illumina) following the Illumina reference guide instructions and recommendations. For each sample, 100 ng of lrPCR mtDNA product was tagmented, purified and ligated to IDT-ILMN UD indexes (Illumina). Samples were pooled by volume using an index correction factor (from Illumina technical note “Balancing sample coverage for whole-genome sequencing and its associated index correction values”). Library pools were quantified with Qubit ssDNA (single-stranded) assay (Invitrogen) and molarity values were calculated considering 450 bp as the average library size.

Paired-end DNA sequencing (2×151 bp) of the libraries was performed on the NovaSeq 6000 sequencer (Illumina) using the NovaSeq S4 Reagent Kit, v1.5; 300 cycles (Illumina) following the manufacturer’s protocol. Images analysis, basecalling and quality scoring of the run were processed using the manufacturer’s software Real Time Analysis (RTA v3.4.4) and followed by generation of FASTQ sequence files.

### Pre-processing Raw Data

At the time of the analysis, the pre-processing protocol of the raw Oxford Nanopore reads produced with the Q20+ ligation sequencing kit was still under development (https://nanoporetech.com/about-us/news/new-nanopore-sequencing-chemistry-developers-hands-set-deliver-q20-99-raw-read), so we used a custom snakemake pipeline^46^ (https://github.com/marcDabad/q20plus_rebasecall), which involves four different tools. First, the basecalling of the simplex data was performed using Guppy v5.0.16 (https://nanoporetech.com/community). Then, the Duplex Sequencing Tools v0.2.3 (https://github.com/nanoporetech/duplex-tools/) was used to identify and filter the paired duplex reads. Following, the putative duplex reads filtered were high quality basecalled using Guppy Duplex Basecalling beta v0.0.0 (https://nanoporetech.com/community). The last step consisted of merging and removing the redundancy from the two basecalling data results. For this task, we applied an in-house developed tool called fastq_merge. Two models to perform the basecalling were applied: res_dna_r9.4.1_e8.1_sup_v033.cfg (from the Rerio Database) for the FLO-MIN106D (R9.4.1) flow cell generated data, and dna_r10.4_e8.1_sup.cfg (from Guppy) for the FLO-MIN112 (R10.4) data. Quality control checks were performed with MinIONQC.R^47^ which uses the final sequencing_summary.txt file.

### Alignment and variant calling

Alignment of nanopore reads for demultiplexing was performed using minimap2 v2.24-r1122 to version GRCh38 of the human genome producing a PAF alignment file. The fastq files were demultiplexed into individual files per cut site using ont_demult v0.2, which takes as input the original fastq files, the PAF alignment file from minimap2 and a configuration file that gives the location of the cut sites and the correspondence between sample identifiers and cut sites. The demultiplexed files were then re-aligned using minimap2, this time to the mitochondrial genome alone (from GRCh38), producing BAM alignment files. The BAM files corresponding to each sample (from different cut sites) were merged to produce the sample BAM files. Analysis of sequencing depth and variant calling were performed using ont_align_view v1.0 which produces a file showing per base coverage, a VCF file with the called variants, and a text file with all read data aligned together with 1 column per base position and 1 line per read (used for frequency estimation of haplotypes).

Short-read data was processed following the Broad Institute’s best practices for variant calling of mitochondrial variants. Briefly, starting from a whole genome alignment, those reads mapping to the chrM were extracted and aligned with BWA-MEM to a reference containing only chrM sequence or chrM shifted 8Kb. Variant calling was performed on both alignments with mutect (GATK v.4.1.7.0) and the variants on the shifted alignment were lifted over to the regular chrM sequence.

### Annotation of mtDNA variants

Variants called were converted to vcf format with bcftools^48^. Heteroplasmy fraction of each variant was computed and appended to the vcf file with a custom script. Briefly, heteroplasmy fraction in a given position was computed by dividing the sum of reads supporting the alternative allele by the total number of reads spanning that position. Sample major and minor haplogroups were determined with Haplocheck^38^.

Variants were annotated with functional annotations, population frequencies and pathogenicity predictors with SnpEff and SnpSift^49,50^. gnomAD, MITOMAP and MitoTIP^35–37^ were used as sources of annotation. Relevant information fields of each variant were extracted and converted to a tabular format with bcftools.

Possible disease-associated variants were prioritized based on heteroplasmic fraction, population frequencies and “Confirmed” or “Reported” status in MITOMAP. Variant filtering was performed with bcftools.

### Circos plot

For creating the plots, we first mapped the reads using the software Minimap2 (2.24). Then we extracted only the reads belonging to the mitochondrial chromosome and we computed the coverage for each locus using Samtools depth (v. 1.14). The output was parsed to create files appropriately formatted for input to Circos (v. 0.69-9).

## Acknowledgements

This research has received funding from the European Union’s Horizon 2020 research and innovation programme under grant agreement No 824110 – EASI-Genomics, the ERC Synergy project BCLL@las under grant agreement No 810287 and from the Spanish Instituto de Salud Carlos III, Fondo de Investigaciones Sanitarias and cofounded with ERDF funds (PI19/01772).

We acknowledge the support of the Spanish Ministry of Science and Innovation through the Instituto de Salud Carlos III and the 2014–2020 Smart Growth Operating Program, to the EMBL partnership and cofinancing with the European Regional Development Fund (MINECO/FEDER, BIO2015-71792-P). We also acknowledge the support of the Centro de Excelencia Severo Ochoa, and the Generalitat de Catalunya through the Departament de Salut, Departament d’Empresa i Coneixement and the CERCA Programme.

## Author contributions

M.G. and I.G.G. conceived the project. I.K., P.B., M.G. and I.G.G. developed the method. I.K. performed experiments. I.B.H. provided cell line samples. A.A. and E.G.A. contributed to patient recruitment and provided clinical samples; J.L. and E.G.A. processed and analyzed the clinical samples at the diagnostic lab. S.H. developed informatics pipeline. S.H., D.C., M.C.F.L., M.D., R.T.H., I.P., M.J.I. processed and analyzed the data. I.K., S.H., M.G. and I.G.G. wrote the manuscript. All authors revised and approved the manuscript for publication.

## Additional information

Correspondence and requests for materials should be addressed to Marta Gut and Ivo Glynne Gut.

