## Supplementary for "Novel method for multiplexed full-length single-molecule sequencing of the human mitochondrial genome"

### (Supplementary Material)

Ieva Keraite[1], Philipp Becker[1‡], Davide Canevazzi[1], Maria C. Frias-López[1], Marc Dabad[1], Raúl Tonda-Hernandez[1], Ida Paramonov[1], Matthew John Ingham[1], Isabelle Brun-Heath[2,3], Jordi Leno[4,5], Anna Abulí[4,5], Elena Garcia-Arumí[4,6,7], Simon Heath[1,8], Marta Gut[1,8*] & Ivo Glynne Gut[1,8*]

[1] CNAG-CRG, Centre for Genomic Regulation (CRG), The Barcelona Institute of Science and Technology (BIST), Barcelona, Spain

[2] Institute for Research in Biomedicine (IRB Barcelona), The Barcelona Institute of Science and Technology (BIST), Barcelona, Spain.

[3] Joint IRB-BSC Program in Computational Biology, Barcelona, Spain

[4] Department of Clinical and Molecular Genetics and Rare Disease, Hospital Universitari Vall d'Hebron, Barcelona, Spain

[5] Medicine Genetics Group, VHIR, Hospital Universitari Vall d'Hebron, Barcelona, Spain

[6] Research Group on Neuromuscular and Mitochondrial Disorders, VHIR, Hospital Universitari Vall d'Hebron, Barcelona, Spain

[7] Centro de Investigación Biomédica en Red de Enfermedades Raras (CIBERER), Instituto de Salud Carlos III, Barcelona, Spain.

[8] Universitat Pompeu Fabra, Barcelona, Spain

[‡] Current address: Qiagen, Hilden, Germany

*All correspondence should be addressed to:

**Supplementary results.**

**Guide RNA (gRNA) selection and multiplexing strategy.** We used the Geneious Prime 2021.2.2 tool to identify 2,198 potential guides, of which 428 were complementary to the light strand (LS) and 1,770 to the heavy strand (HS). Once filtered according to the Doench (2014) Activity Score[1] and disregarding all sequences that have a score of <0.3, we obtained 132 LS and 397 HS guide sequences (data not shown). This classification only informs on the level of cleavage efficiency based on the observations of guide RNA sequence features. It is speculated that there are differences between biochemical and cell-based Cas9 cleavage due to RNA folding, stability, complex formation[2], and available off-target sequences. In our approach for designing synthetic CRISPR RNA (crRNA) sequences the most important factor was a low number of off-targets resulting in stringent enrichment of mtDNA molecules (avoiding hypervariable regions and well-described variants). We generally used two guides in two separate aliquots for each sample, as shown in Fig. 1. The guides are located approximatelly 8 kb from each other and one targets the HS and the other the LS, in order to produce high coverage data over the cut sites and to identify areas where cutting efficiency is reduced due to mtDNA alterations. In more complex situations, when analysing deletions, it is possible to use more than two guides in a corresponding number of aliquots per sample located in strategically selected positions. We desgined and tested four gRNA pairs spanning the entire mtDNA sequence considering hypervariable regions and major pathogenic variants, to multiplex up to four samples (Supplementary Table 1).

**Implementation and optimization of Cas9-mtDNA-enrichment**. First, to validate the method and to demonstrate the preferential sequencing of the full-length mtDNA molecule with high output we used a gDNA sample in two aliquots with one pair of gRNAs. Here we used high integrity gDNA extracted from the human embryonic kidney HEK293 cell line to enrich for intact full-length mtDNA and to demonstrate reliable demultiplexing. As a control we added three blank (without gDNA) samples incubated each with a different pair of gRNAs. We achieved a coverage of the targeted region exceeding 5,000× (Supplementary Table 4). We also observed an extremely low number of reads (<0.02%) with non-intended cuts from the blank reactions, confirming our approach for reliable demultiplexing of multiple pooled samples (Supplementary Table 4). Next, we focused on increasing the data output. It was indicated that the Cas9 complex stays bound to the 5' end of the guide RNA[3]. In our case meaning the Cas9 complex stays bound to the end of the linearized mtDNA molecules. This may obstruct transition of the DNA molecule through the nanopore and reduce sequencing yield[4]. Therefore we used Proteinase K to digest the Cas9 nuclease. By introducing the Proteinase K digest, we achieved a 2-fold increase of the full-length reads fraction mapping to the mtDNA reference.

To reduce the nuclear DNA fraction in the starting material and to increase the final enrichment of the circular mtDNA molecules we introduced an Exonuclease V treatment upstream. In this strategy we used high integrity gDNA extracted from HEK293, CAPAN-2, A549, SH-SY5Y cell lines and we achieved a mean increase of the mtDNA fraction from 27% to 49% of total reads on a GridIon flow cell (Supplementary Table 5). Performing DNA quality control upfront the gDNA processing is very important to decide whether introducing Exonuclease V treatment step will be useful. Inclusion of the Exonuclease V treatment step for low integrity gDNA samples is counterproductive as it leads to digestion of fragmented mtDNA (data not shown).

**Multiplexed sequencing assay with different GridION flow cells**. We applied our method to four cell line DNA samples (HEK293, CAPAN-2, A549, SH-SY5Y) utilizing four pairs of gRNAs for multiplexing, Proteinase K and Exonuclease V. The library of the pooled samples was prepared with the Q20+ ligation sequencing kit (Kit 12 Chemistry) and run on different GridION flow cell versions - R9.4.1, R10.3 and R10.4 (Supplementary Table 5). The percentage of on-target reads mapping to mtDNA reached >60% on R9.4.1 flow cells, 57% on

R10.3 and 52% on R10.4 flow cells. The coverage of the full-length mtDNA in the library pool ranged from 4,500× to 11,559× on the R9.4.1 flow cell vs. 2,002× and 5,040× on the R10.3 flow cell after demultiplexing. Comparing the pooled library on R9.4.1 vs R10.4, coverage ranged from 1,739× to 8,318× vs. 2,347× to 9,444×, respectively. In this comparison, the lowest output as expected was in the R10.3 flow cell, while the yield obtained with R9.4.1 and R10.4 flow cells was comparable.

**Improving average divergence per base.** Until recently, nanopore sequencing showed a high error rate of up to 10%. The Q20+ ligation sequencing kit together with the improved double-reader head R10.4 pore flow cells were introduced to ameliorate the single molecule read accuracy, especially in complex genomic regions with homopolymers. Originally, we used the ligation sequencing kit SQK-LSK110 and measured the average divergence per base at 4.15–5.58% (data not shown). Changing to the Kit 12 Chemistry resulted in eminent improvement across all types of GridION flow cells currently available from ONT (Supplementary Table 6). The average divergence per base measured was 1.07 ± 0.03% on R9.4.1 and 1.05 ± 0.04% on R10.3 ($P$ = 0.3404), while R10.4 demonstrated a significant improvement in comparison to R9.4.1 flow cell, 1.00 ± 0.04% vs. 1.08 ± 0.03% ($P$ = 0.0043), respectively.

**Validation of the variant calling pipeline and assessment of multiplexing bleed-through.** To validate the multiplexing strategy reliability we compared the nanopore sequencing genotype data to Illumina short-read sequencing of mtDNA enriched by long-range PCR (lrPCR) in gDNA samples from the four cell lines above. The GATK pipeline was used to call variants from the Illumina data of the four cell line gDNA samples. Across the mtDNA genome, 44 positions were identified by GATK where at least one sample had an alternative allele in homoplasmy (Supplementary Table 7a). The same 44 positions were also identified as homoplasmic variants from the ONT sequencing data, and the variant calls were completely concordant across all four flow cells as well as with the Illumina short-read sequencing results.

Additionally, 26 positions were called from the Illumina data as being heteroplasmic with estimated alternate allele frequencies >= 0.5%; none of the putative heteroplasmic variants were shared across multiple samples (Supplementary Table 7b). All 26 sites were also identified (with the same alternate allele) using the ONT data, although for some sites the variant call was indicated as being of low quality with one or more of the flow cells. Estimates of alternate allele frequencies from Illumina and ONT were highly concordant (Supplementary Fig. 1). Sensitivity and specificity were calculated for the 4 flow cells considering variants called with frequency >= 0.5% and treating the called variants from the Illumina data as the truth. The estimated specificity for all 4 flow cells was >99.9%, and the sensitivity was 98.8% for the R10.3 flow cell and 97.7% for the R10.4 flow cell, and slightly lower at 94.2% and 96.5% for the two R9.4.1 flow cells. A blacklist of mtDNA reference sequence positions was established for automatic filtering from the variant calling pipeline (Supplementary Table 7c).

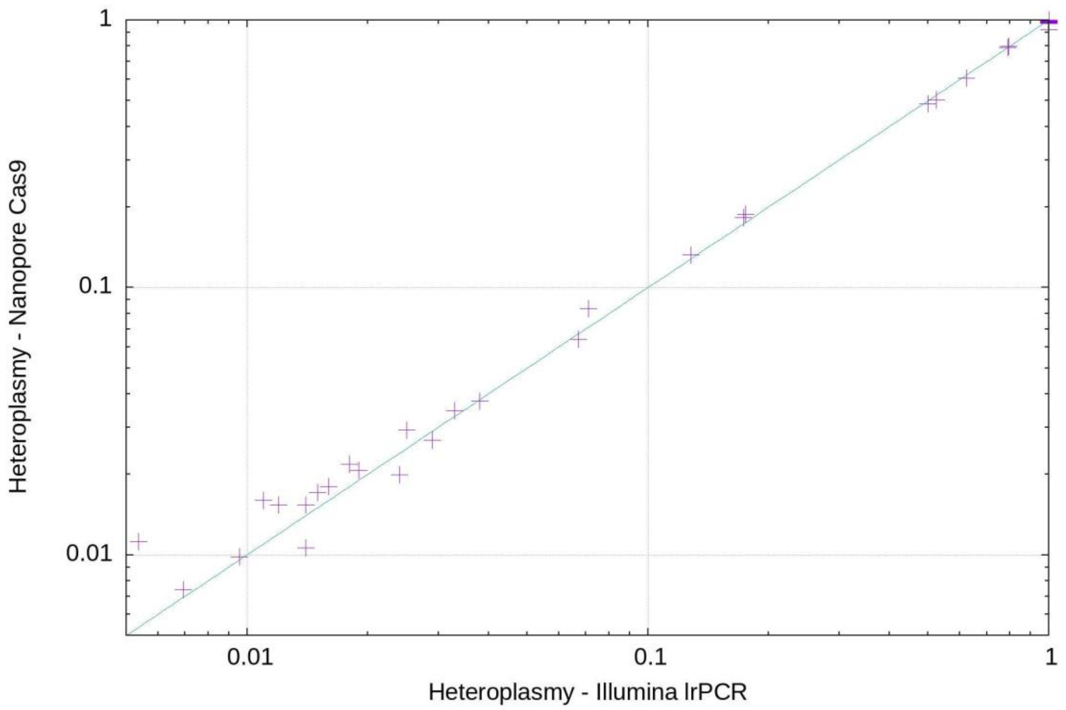


**Supplementary Fig. 1 | Correlation between the mtDNA heteroplasmy frequencies in cell lines measured by lrPCR Illumina and Cas9-mtDNA-enrichment ONT sequencing.**

Three high frequency heteroplasmic variants were identified in the SH-SY5Y cell line from both the Illumina and ONT data (m.1217G>A, m.15490C>A, m.15961G>T) with estimated frequencies from ONT of 19.2%, 17.8% and 49.2% respectively. Using the full-length mtDNA reads, we are able to determine all of the haplotypes of the 3 variants present in this sample. Out of the 8 possible haplotypes, GCT, GCG and AAG are present at high frequency (48.6%, 32.0% and 18.4% respectively) with the other 5 haplotypes making up the remaining 1%. In short-read Illumina sequencing we confirmed the SNVs frequencies but it was only possible to resolve the phase of the two variants positioned nearby each other (m.15490, m.15961), due to the inherent limitations of short-read sequencing (Supplementary Fig. 2).

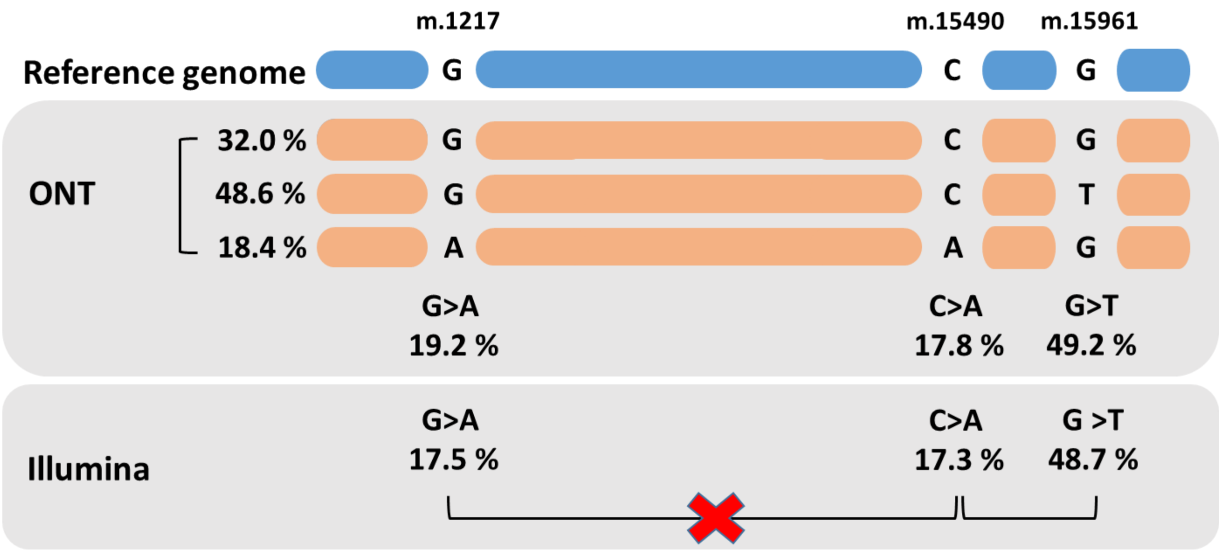


**Supplementary Fig. 2 | mtDNA SNVs phasing with long- and short-read sequencing.** Visual representation of resolved SH-SY5Y cell line mtDNA haplotypes of the distant variants by nanopore sequencing. Using Illumina data only, the m.1217-m.15490 phased SNVs stay unresolved.

**Quality control of clinical samples by lrPCR.** Full-length mtDNA amplification product showed SVs in one clinical sample AW6506. We observed two bands representing two mtDNA populations of ~13.5 kb and ~3.5 kb in this sample. Wild-type 16.5 kb mtDNA was not visible on the gel, due to the amplification bias against larger fragments (Supplementary Fig. 3).

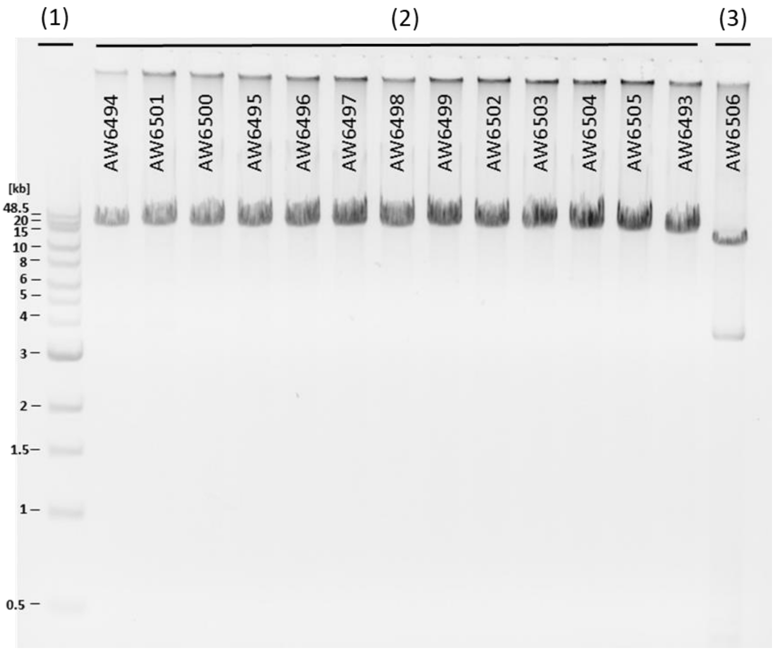


**Supplementary Fig. 3│Agarose gel of the lrPCR products from the clinical samples.** (1) molecular weight marker, (2) amplicons with 16.5 kb mtDNA and (3) amplicons with two mtDNA SVs in sample AW6506.
